# Thermal plasticity of wing spot size in *Drosophila guttifera*: investigating the relevance to Wingless morphogen

**DOI:** 10.1101/2023.01.09.523128

**Authors:** Yuichi Fukutomi, Aya Takahashi, Shigeyuki Koshikawa

**Affiliations:** Department of Evolution and Ecology, University of California, Davis, One Shields Ave, Davis, CA, 95616, USA; Department of Biological Sciences, Tokyo Metropolitan University, 1-1 Minamiosawa, Hachioji 192-0397, Japan; Research Center for Genomics and Bioinformatics, Tokyo Metropolitan University, 1-1 Minamiosawa, Hachioji 192-0397, Japan; Graduate School of Environmental Science, Hokkaido University, N10W5, Kita-ku, Sapporo, Hokkaido 060-0810, Japan; Faculty of Environmental Earth Science, Hokkaido University, N10W5, Kita-ku, Sapporo, Hokkaido 060-0810, Japan

**Keywords:** *Drosophila guttifera*, thermal plasticity, color pattern formation, morphogen, image binarization

## Abstract

Thermal plasticity of animal color patterns has been studied to investigate the molecular mechanisms of phenotypic plasticity. There are examples in which the formation of color patterns is controlled by a morphogen. The extracellular distribution of a morphogen can be plastic with temperature change. Whether alteration of morphogen distribution with temperature change produces thermal plasticity in color patterns has not been studied. To address this question, polka-dotted wing spots of *Drosophila guttifera*, whose inducer is *wingless* morphogen, can be a suitable model system. In this study, we reared *D. guttifera* at different temperatures to test whether wing spots show thermal plasticity. We found that wing size becomes larger and spot size adjusted with wing size becomes smaller at lower temperatures. We also changed the rearing temperature in the middle of the pupal period and found that the most sensitive developmental period for wing size and spot size is different. The results suggest that developmental mechanisms for the thermal plasticity of wing size and spot size are different. We also found that the most sensitive stage for spot size was part of the pupal period including stages at which *wingless* is expressed in the polka-dotted pattern. Therefore, it is suggested that temperature change might affect the process of specifying spot areas by Wingless in the extracellular region.

## 1 Introduction

Organisms show changes in morphology, physiology, or behavior when they face different environmental conditions and these phenomena are called phenotypic plasticity (West-Eberhard 1989). Phenotypic plasticity of animal color patterns can be observed in various taxa, for example, thermal plasticity of pigmentation patterns on the wings of butterflies and the abdomen of fruit flies, changes of feather coloration on in response to diet, and rapid camouflage of flatfishes to match their backgrounds et al. 1990; Ramachandran et al. 1996; Price 2006; Lafuente et al. 2021). Investigation developmental mechanisms for color pattern formation has contributed greatly to the understanding of molecular mechanisms for phenotypic plasticity of animal color patterns (Tschirren et al. 2003; Gibert et al. 2007; De Castro et al. 2018; van der Burg et 2020).

As mechanisms for the formation of animal color patterns, theoretical models predict that diffusible factors such as morphogens (signaling molecules that determine patterns of animal body plans in ontogeny; Wolpert 1969) specify those patterns (Turing 1952; Murray 1981; Kondo and Shirota 2009). As empirical examples, there are cases that diffusible signaling molecules control or are considered to control the formation of color patterns such as wing pigmentation patterns of butterflies and a fruit fly species, *Drosophila guttifera* (Werner et al. 2010; Martin et al. 2012; Mazo-Vargas et al. 2017; Özsu et al. 2017). Among those cases, wing color patterns of *Junonia coenia* and *Bicyclus anynana* are known to show thermal plasticity. *WntA* gene is responsible for the formation of some components of the color pattern on the wings of *J. coenia* (Mazo-Vargas et al. 2017) and the color pattern exhibits plasticity upon cold shock (Nijhout 1984; Serfas and Carroll 2005). *wingless* gene is considered to specify the pattern of eyespots on wings of *B. anynana* (Özsu et al. 2017) and the size of the eyespot changes when the butterflies are reared at different temperatures (Brakefield et al. 1998). The extracellular distribution of diffusible signaling molecules such as morphogens is considered to be plastic to the surrounding temperature (Eldar et al. 2004; Barkai and Shilo 2009).

How the change in temperature affects the distribution of a morphogen is studied in embryos of *Drosophila melanogaster*. It is shown that the distribution of Bicoid morphogen, which specifies the anteroposterior axis, exhibits a drastic change when reared under different temperature conditions (Houchmandzadeh et al. 2002). In the context of color pattern formation in wings, it is considered that changes in the distribution of signaling molecules such as WntA and Wingless will alter the outcome patterning (Martin and Reed 2014; Martin and Courtier-Orgogozo 2017). However, how temperature changes affect the distribution of signaling molecules responsible for the formation of color patterns is totally unknown.

*Drosophila guttifera* has a polka-dotted melanin pigmentation pattern on the wing. The color pattern of this species has been used in the context of multiple research fields (Fukutomi and Koshikawa 2021; Niida and Koshikawa 2021). In this species, melanin spots can be observed around campaniform sensilla (Lees 1942) and *wingless* gene is expressed at the campaniform sensilla during the pupal period (Werner et al. 2010; Koshikawa et al. 2015; Koseki et al. 2021). Wingless is an inducer of wing pigmentation and is assumed to specify the area of wing spots by diffusion from campaniform sensilla (Werner et al. 2010). *D. guttifera* can be a suitable model to investigate how temperature change affects color pattern formation by a diffusible factor, but whether wing spots of *D. guttifera* show thermal plasticity has not been investigated. In this study, we reared *D. guttifera* at different temperatures, and measured wing size and spot size. We found that wing size, spot size itself, and spot size adjusted with wing size shows thermal plasticity. We also found that part of the pupal period including stages at which *wingless* is expressed in the polka-dotted pattern is the most sensitive period for thermal plasticity.

## 2 Materials and methods

### 2.1 Rearing flies and preparing samples

A fly stock used in this study was a wild-type strain of *D. guttifera* (stock no. 15130-1971.10) from *Drosophila* species stock center at the University of California, San Diego. Flies were reared with malt food (containing 50 g cornmeal, 50 g malt, 50 g sugar, 40 g yeast, and 5 g agar in 1 litter of water). For experiments, 10 adult male flies and 10 adult female flies were crossed in one vial. Adult flies for crosses were removed four days later. When progenies became adults, they were collected within 24 hours after eclosion. To analyze the phenotypic plasticity of wing spots, we reared flies under 18 °C, 21 °C, 25 °C, and 28 °C. As food tends to be dried up at higher temperatures, we placed all fly vials in plastic bags in which moist tissue paper is placed.

For experiments to change the temperature during pupal stages, we picked up pupae at stage P4 (i) (Fukutomi et al. 2017; Fukutomi et al. 2018) and moved them onto moist tissue paper in a Petri dish. P4 (i) is the distinguishable stage without dissection and it is right before the stage when the expression of *wingless* starts in the polka-dotted pattern (Werner et al. 2010; Fukutomi et al. 2017). Flies were reared under the following three conditions.

“Condition 1”:

Until P4 (i), flies were reared at 18 °C. From P4 (i), they were reared at 25 °C.

“Condition 2”:

Until P4 (i), flies were reared at 25 °C. From P4(i) to P14-15 (stages of pupae), they were reared at 18 °C. At P14-P15, they were moved back to an incubator at 25 °C.

“Condition 3”:

Until P14-15, flies were reared at 25 °C. From P14-15, they were reared at 18 °C.

Six to seven days after eclosion, flies were anesthetized with CO_2_. Right wings were dissected and mounted. For mounting solution, a mixture of Hoyer’s solution and lactic acid (1:1 ratio) was used. Photo images were taken with a digital camera (DP73, OLYMPUS) connected to a microscope (SZX16, OLYMPUS). For the backgrounds of photo images, the reference greyscale (Brightness = 128, ColorChecker, Xrite) was used.

### 2.2 Measuring and analyzing wing size and spot size

At first, the brightness of the backgrounds of all photo images was adjusted with ImageJ software (Schneider et al. 2012). The upper right region of each photo image was selected by a rectangle (400 x 300 pixels) and the background was converted by “Window/Lebel…” function so that the brightness of the selected region became 128.

To estimate the wing size of *Drosophila*, centroid size is often used (Debat et al. 2003; Abbott et al. 2010). In this study, we calculated the centroid size for each wing by the following procedures. The intersection points of veins (Figure 1a) were used as landmarks for calculating centroid size. As the coordinates of landmarks and centroid of a selected polygon (Figure 1a) in each photo image were provided by ImageJ, the centroid size can be calculated.

**Figure 1.**
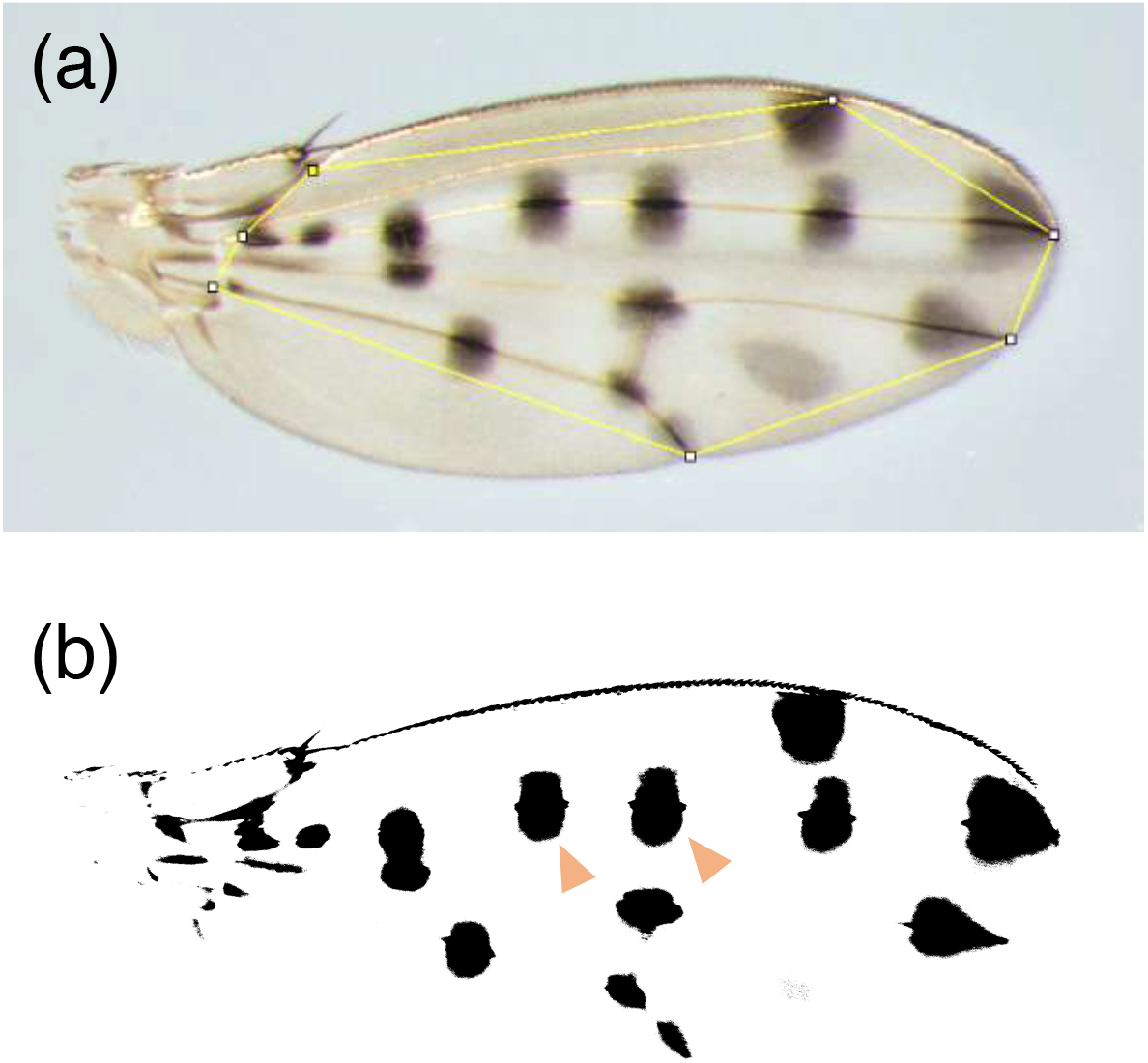
Landmarks and spots on a wing used in this study. (a) Landmarks and a polygon used for analyses. Landmarks, intersection points of veins, are indicated as white dots. The polygon was drawn by connecting white dots. The brightness of the background was increased with ImageJ. (b) Binarized image of a wing. For convenience, we call the spot indicated with the left soft orange arrowhead as “Proximal”, and call the one indicated with the right soft orange arrowhead as “Middle”.

For measuring the spot size, the photo images were binarized with ImageJ. We converted photo images to 8-bit images and selected the polygon indicated in Figure 1a again. Using “Threshold…” function, 8-bit images of wings were binarized and wing spots became black regions (Figure 1b). We adopted Otsu’s method (Otsu 1979) for binarization. By selecting a black region in the binarized images, the area of a wing spot was calculated. The areas of two spots around campaniform sensilla, “Proximal” and “Middle” (Figure 1b), were used for analysis. To adjust the spot size with wing size, spot size was divided by the area of the selected polygon (Figure 1a) instead of centroid size. This is because units of spot size (area, pixels) and centroid size (length, square root of pixels) are different.

Statistical analyses were conducted with R version 4.2.1 (R Core Team 2022). For analyses of wing size and spot size (non-adjusted), one-way ANOVA and Tukey’s honest significant differences (HSD) test were performed. To analyze spot size (adjusted with wing size), we performed Kruskal-Wallis rank sum test and pairwise comparisons using Wilcoxon rank sum test. We used Bonferroni correction for adjustment of *p*-values in pairwise comparisons using Wilcoxon rank sum test. All graphs were produced with ggplot2 (Wickham 2009).

## 3 Results

### 3.1 Wing size and spot size under different temperatures

When flies were reared under four different temperatures, wings appeared to be larger at lower temperatures (Figure 2). We calculated the centroid size and found that it became smaller as the rearing temperature was increased (Figure 3). That tendency was observed both in males and in females with one exception of a nonsignificant difference between the centroid size at 18 °C and 21 °C in males (Figure 3a).

**Figure 2.**
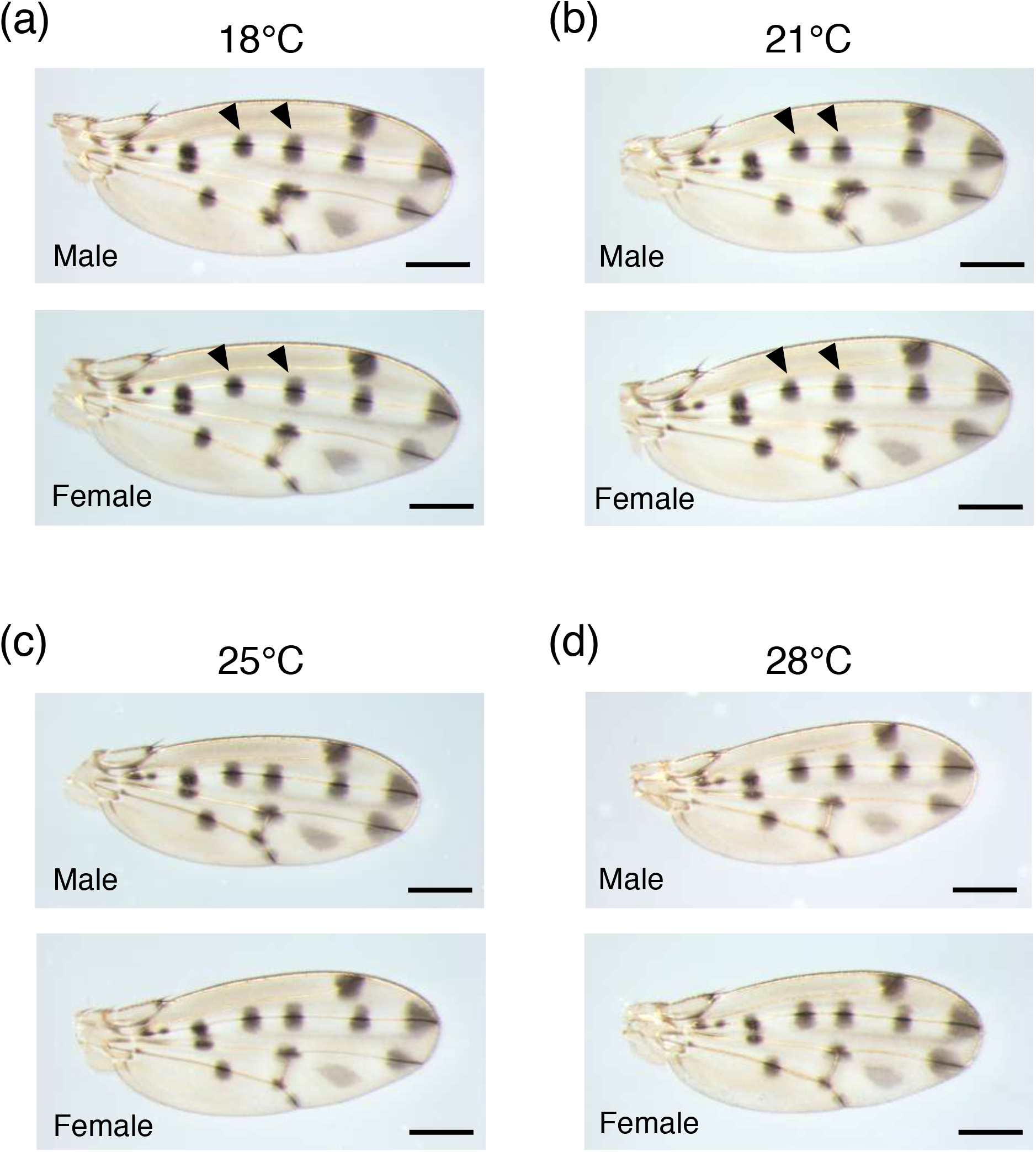
Wings from male and female flies reared at 18 °C, 21 °C, 25 °C, and 28 °C. (a) Wings from flies at 18 °C. The left black arrowheads indicate “Proximal” spots and the right black arrowheads indicate “Middle” spots. (b) Wings from flies at 21 C. (c) Wings from flies at 25 C. (d) Wings from flies at 28 °C. For all pictures, the brightness of the background was increased with ImageJ. Scale bars indicate 400 μm.

**Figure 3.**
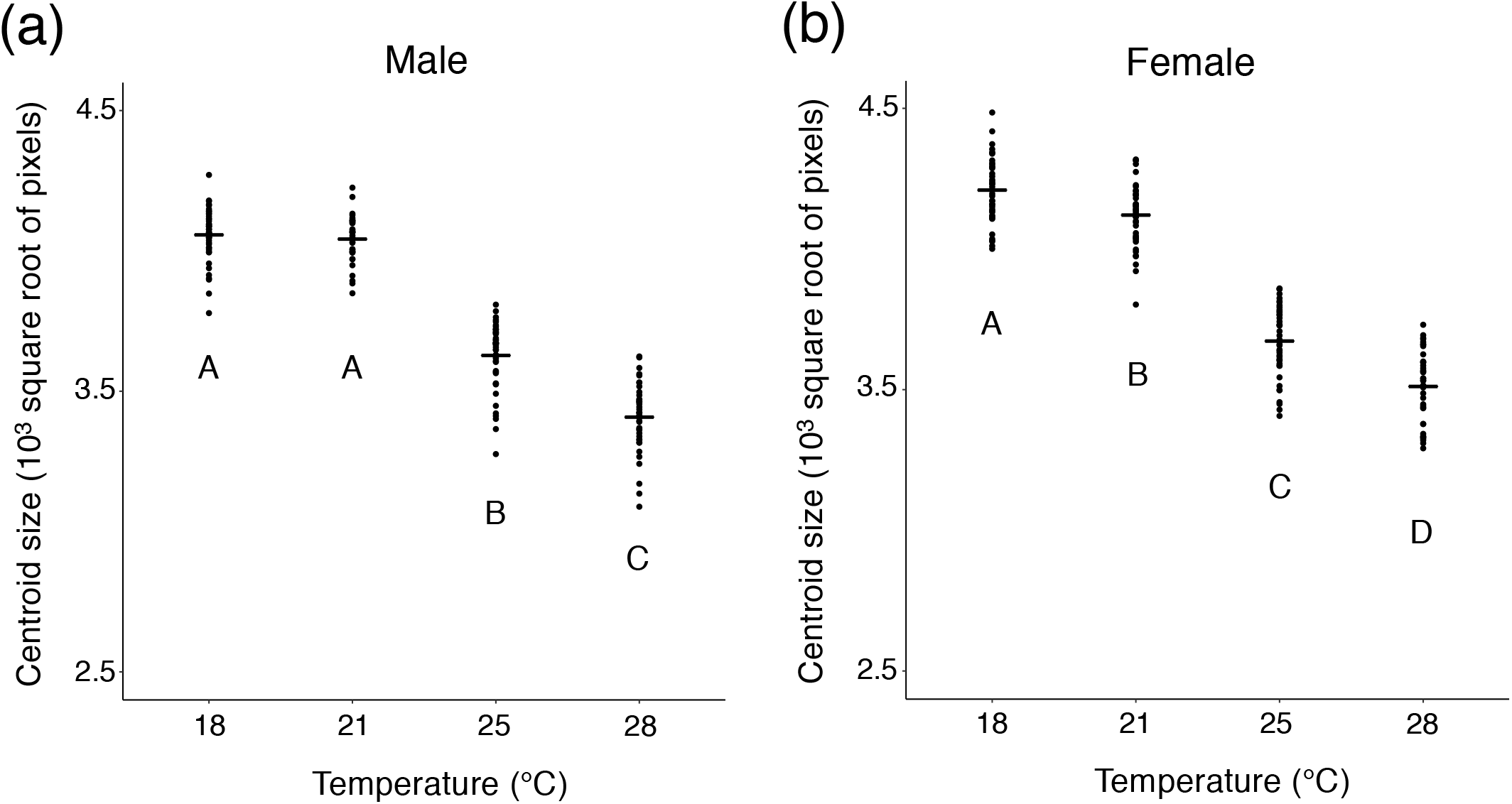
Centroid size of wings from flies reared at 18 °C, 21 °C, 25 °C, and 28 °C. (a) Centroid size of wings from male flies. (b) Centroid size of wings from female flies. In males and females, there were significant differences between temperatures(*p* < 10^-15^, one-way ANOVA, degree of freedom = 3, *F* = 383.9 in (a), 331.1 in (b)). Different letters indicate significant differences (*p* < 0.05, Tukey’s HSD test). Black bars indicate mean values.

We measured the spot size and found that there was no clear tendency correlated with temperature although significant differences between different temperature were detected in all categories by one-way ANOVA (Figure 4). The size of “Proximal” spots was almost stable in male flies, as no significant difference was detected between the spot size at 18 °C, 25 °C, and 28 °C by Tukey’s HSD test (Figure 4a). The size of “Proximal” spots in female flies was not as stable as that in male flies (Figure 4b). It is possible to interpret that the size of “Middle” spots in males became smaller when the temperature got higher (Figure 4c), but it is not possible to interpret the data on the size of “Middle” spots in females in the same way (Figure 4d).

**Figure 4.**
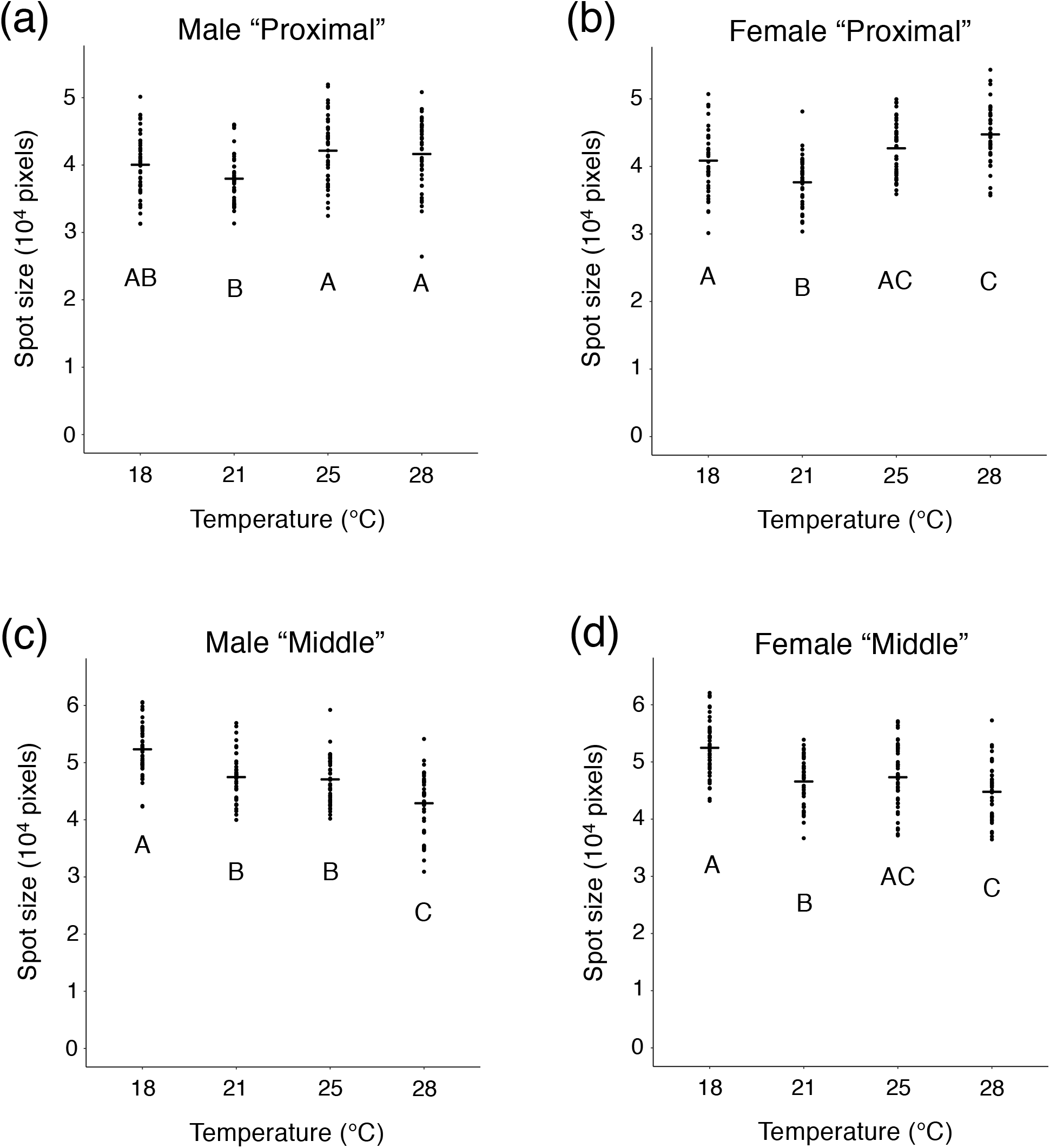
The spot size of wings from flies reared at 18 °C, 21 °C, 25 °C, and 28 °C. (a) Size of “Proximal” spots on wings from male flies. (b) Size of “Proximal” spots on wings from female flies. (c) Size of “Middle” spots on wings from male flies. (d) Size of “Middle” spots on wings from female flies. In all categories, there were significant differences between temperatures (*p* < 0.001, one-way ANOVA, degree of freedom = 3, *F* = 6.546 in (a), 20.83 in (b), 31.25 in (c), 18.6 in (d)). Different letters indicate significant differences (*p* < 0.05, Tukey’s HSD test). Black bars indicate mean values.

As centroid size of wings and the area of the polygon mentioned above are highly correlated (Figure 5), we divided the spot size by the area of the polygon to adjust the spot size by the wing size. After the adjustment, we found that the ratio of the spot size to the wing size was higher at higher temperatures (Figure 6). Both in males and females, a significant difference between the ratio of “Proximal” size to wing size at 25 °C and the ratio at 28 °C was observed by Wilcoxon rank sum test with Bonferroni correction (Figure 6a, b). For “Middle” spots, a significant difference between the ratio at 18 °C and that at 21 °C was detected by Wilcoxon rank sum test with Bonferroni correction (Figure 6c, d).

**Figure 5.**
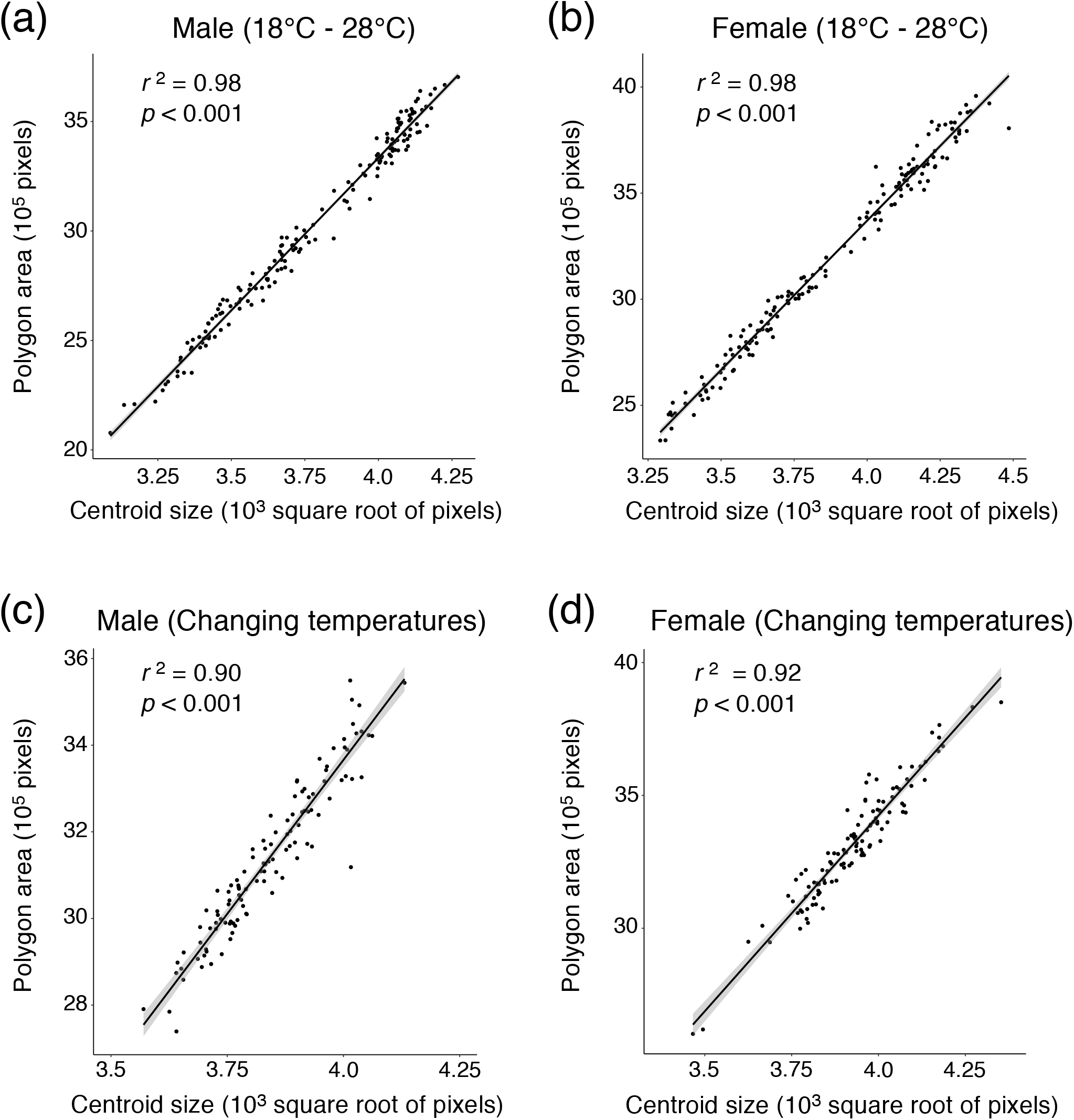
The correlation between centroid size and the area of the polygon. (a) Wings of males reared under 18 °C, 21 °C, 25 °C, and 28 °C. (b) Wings of females reared under 18 °C, 21 °C, 25 °C, and 28 °C. (c) Wings of males whose rearing temperatures were changed during the pupal period. (d) Wings of males whose rearing temperatures were changed during the pupal period. Grey shadows indicate 95% confidence intervals.

**Figure 6.**
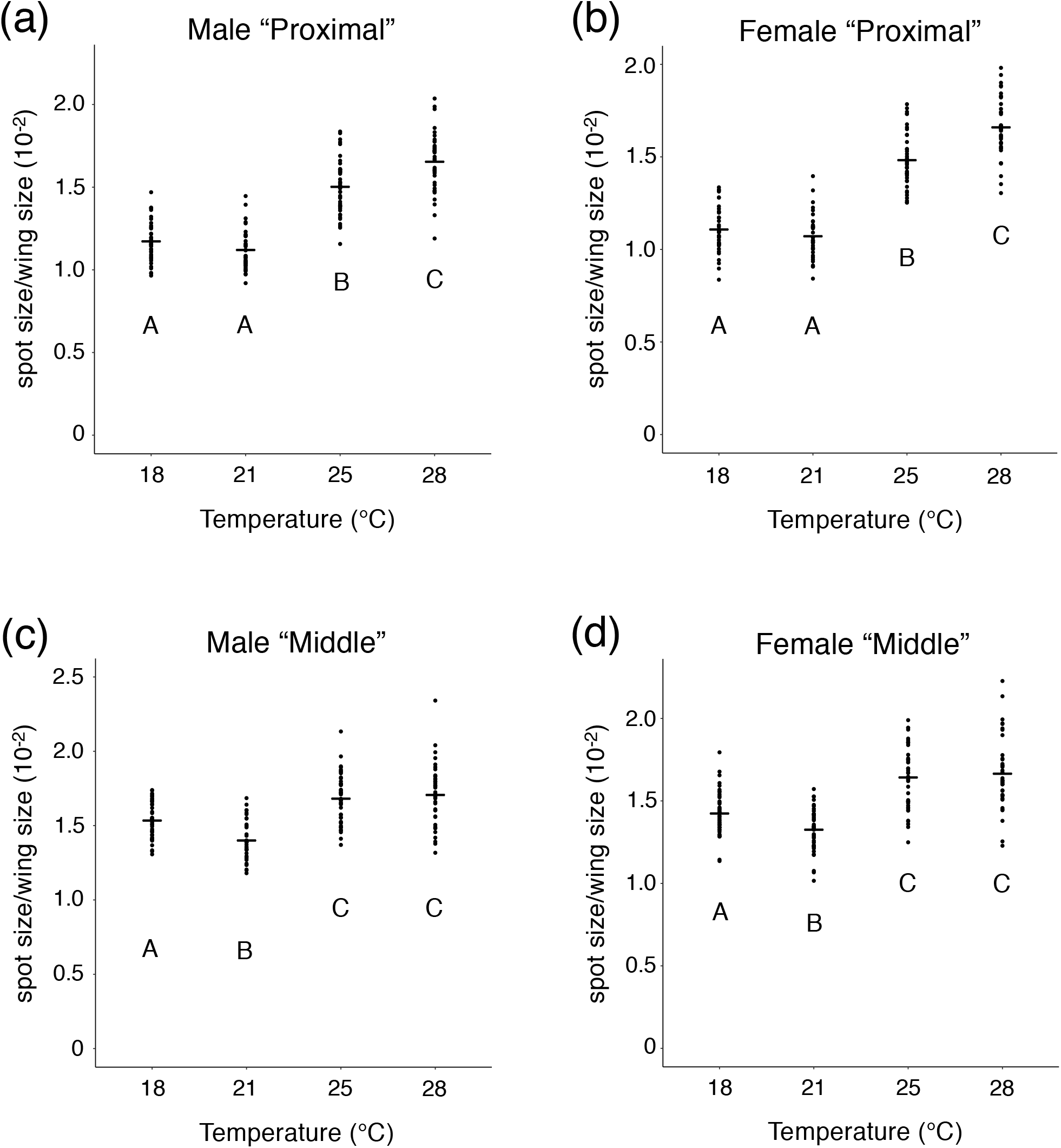
The spot size adjusted with wing size. Flies were reared at 18 °C, 21 °C, 25 °C, and 28 °C. (a) Size of “Proximal” spots adjusted with wing size from male flies. (b) Size of “Proximal” spots adjusted with wing size from female flies. (c) Size of “Middle” spots adjusted with wing size from male flies. (d) Size of “Middle” spots adjusted with wing size from female flies. In all categories, there were significant differences between temperatures (*p* < 10^-13^, Kruskal-Wallis rank sum test, degree of freedom = 3, *χ^2^* = 120.35 in (a), 127.11 in (b), 65.207 in (c), 77.059 in (d)). Different letters indicate significant differences (*p* < 0.05, Wilcoxon rank sum test with Bonferroni correction). Black bars indicate mean values.

As a conspicuous result, we noticed that “Proximal” spot was smaller than “Middle” spot at 18 °C and 21 °C (Figure 2a, b). When we analyzed the ratio of “Proximal” size to “Middle” size, we found that the ratio became higher when the rearing temperature became higher (Figure 7). Other than the comparison between ratios at 18 °C and those at 21 °C, significant differences were detected by Wilcoxon rank sum test with Bonferroni correction both in males and females (Figure 7).

**Figure 7.**
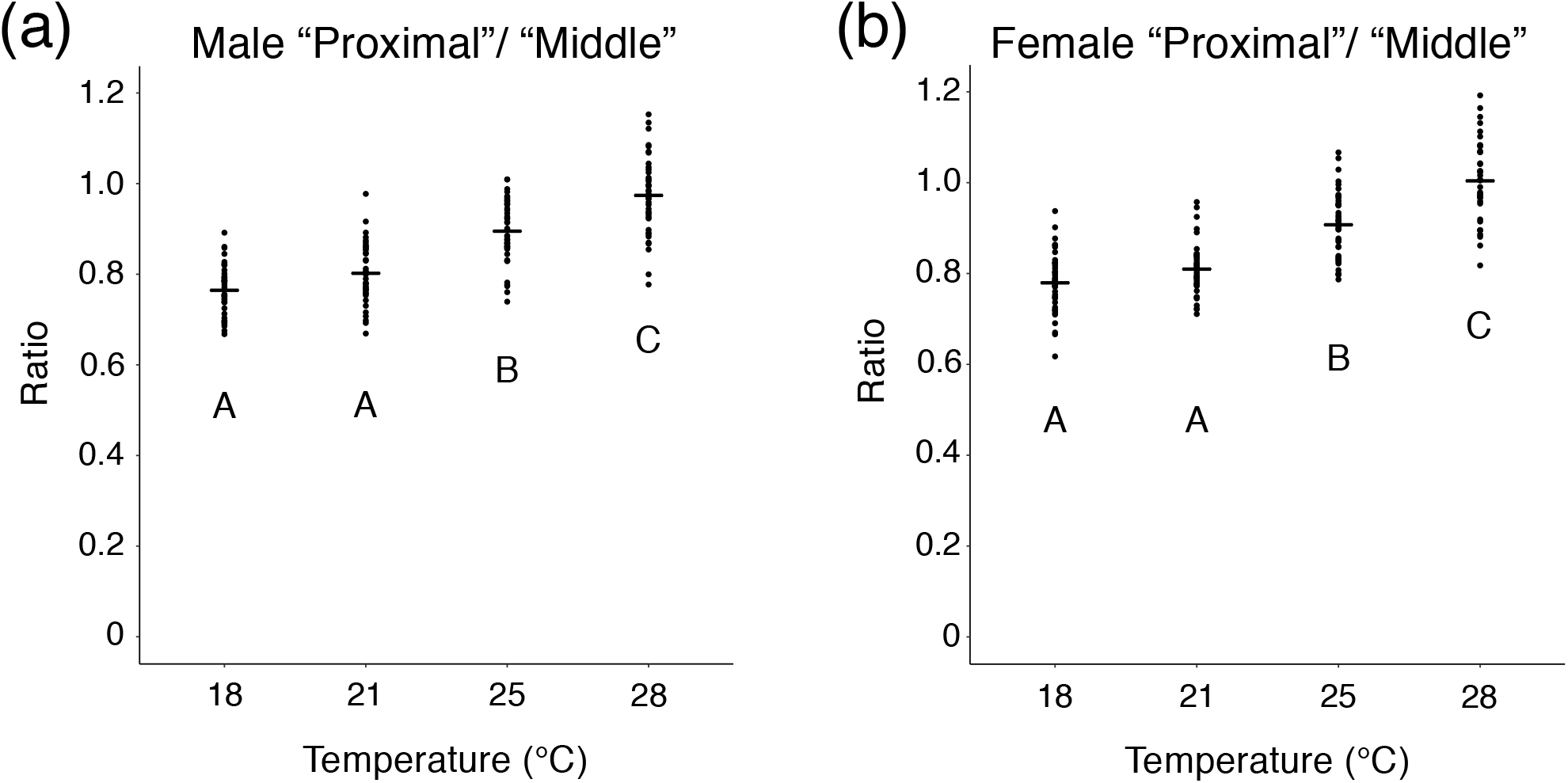
The ratio of “Proximal” spot size to “Middle” spot size. (a) The ratio in male flies. (b) The ratio in female flies. Both in males and females, there were significant differences between temperatures (*p* < 10^-15^, Kruskal-Wallis rank sum test, degree of freedom = 3, *χ^2^* = 101.22 in (a), 102.96 in (b)). Different letters indicate significant differences (*p* < 0.05, Wilcoxon rank sum test with Bonferroni correction). Black bars indicate mean values.

### 3.2 Change of wing size and spot size when the rearing temperature is changed

By rearing *D. guttifera* under different temperatures, it was shown that the wing size and the spot size of *D. guttifera* exhibits phenotypic plasticity. To investigate which stage is sensitive to temperature, we changed the rearing temperature during the pupal period. The wing size was the largest when flies were reared under “Condition 1” (reared at 18 °C until P4 (i)) (Figure 8). By Tukey’s HSD test, significant differences were detected between the wing size of the flies reared under “Condition 1” and the other two conditions (Figure 9). No significant difference was detected between “Condition 2” and “Condition 3” (Figure 9). The same tendency was observed both in males and females (Figure 9a, b).

**Figure 8.**
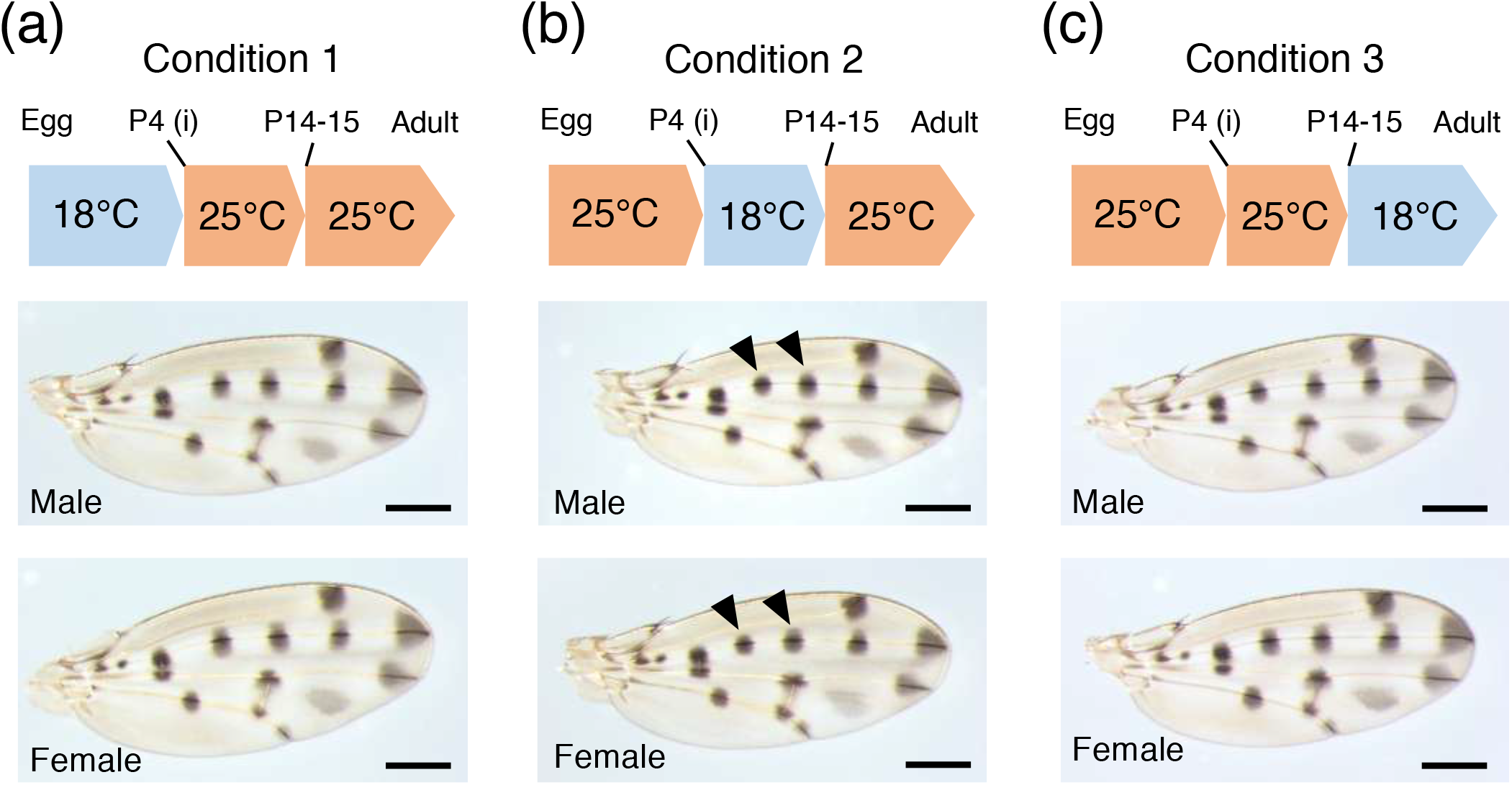
Wings from male and female flies whose rearing temperatures were changed during the pupal period. (a) Wings from male and female flies reared under “Condition 1”. Until P4 (i), flies were reared at 18 °C. From P4 (i), they were reared at 25 °C. (b) Wings from male and female flies reared under “Condition 2”. Until P4 (i), flies were reared at 25 °C. From P4(i) to P14-15, they were reared at 18 °C. From P14-P15, they were reared at 25 °C. The left black arrowheads indicate “Proximal” spots and the right black arrowheads indicate “Middle” spots. (c) Wings from male and female flies reared under “Condition 3”. Until P14-15, flies were reared at 25 °C. From P14-15, they were reared at 18 °C. For all pictures, the brightness of the background was increased with ImageJ. Scale bars indicate 400 μm.

**Figure 9.**
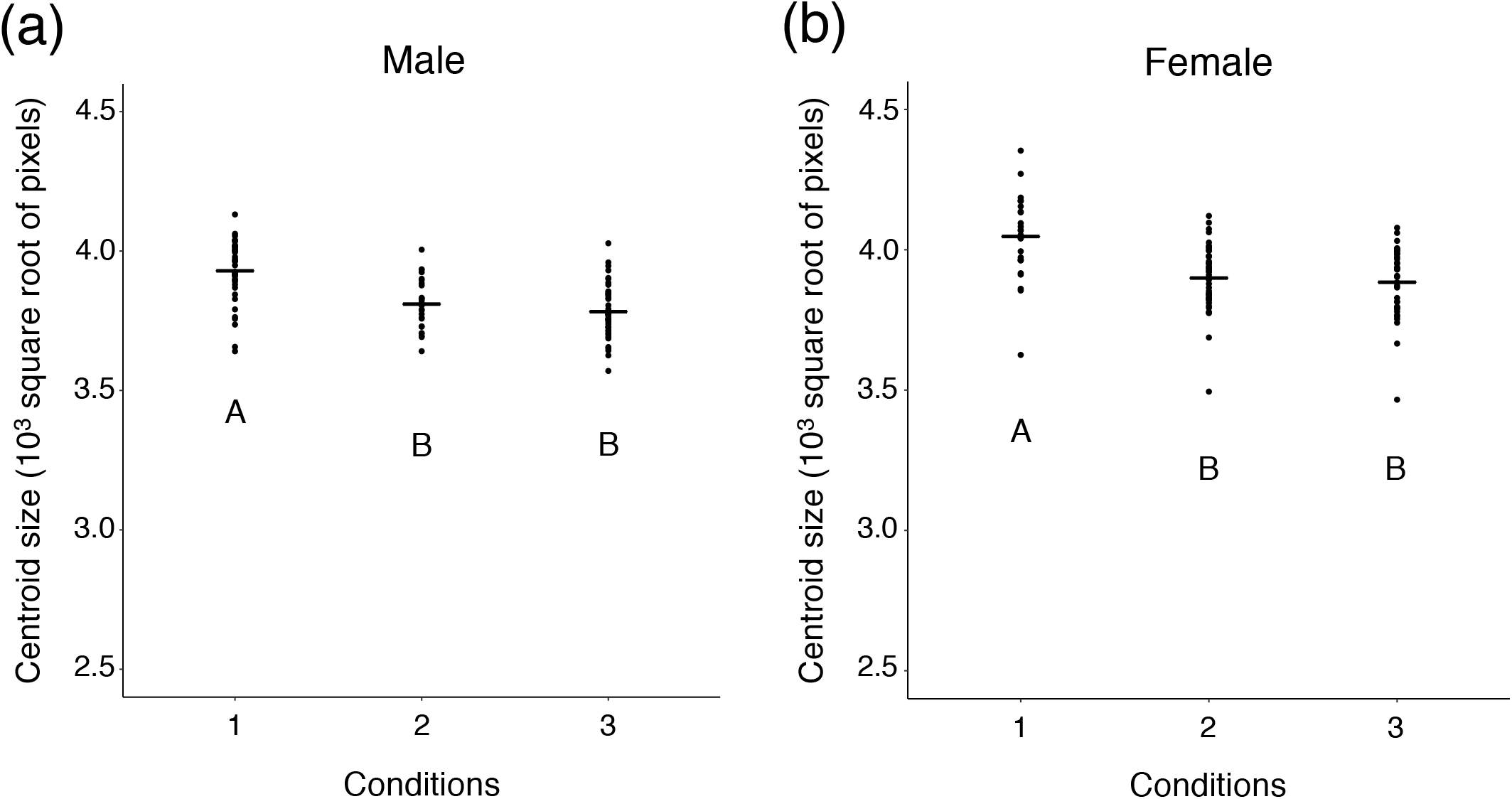
Centroid size of wings from flies reared under “Condition 1”, “Condition 2”, and “Condition 3”. (a) Centroid size of wings from male flies. (b) Centroid size of wings from female flies. Both in males and females, there were significant differences between temperatures(*p* < 10^-5^, one-way ANOVA, degree of freedom = 2, *F* = 24.64 in (a), 15.43 in (b)). Different letters indicate significant differences (*p* < 0.05, Tukey’s HSD test). Black bars indicate mean values.

When we measured spot size, we found that the mean spot size of wings from flies reared under “Condition 2” (reared at 18 °C from P4 (i) to P14-15) became the smallest (Figure 10). Both in males and females, all comparisons of “Proximal” spot size showed significant differences by Tukey’s HSD test (Figure 10a, b). For “Middle” spot size, a significant difference between the spot size of the flies reared under “Condition 1” and “Condition 3” was detected in males by Tukey’s HSD test (Figure 10c), but it was not detected in females (Figure 10d).

**Figure 10.**
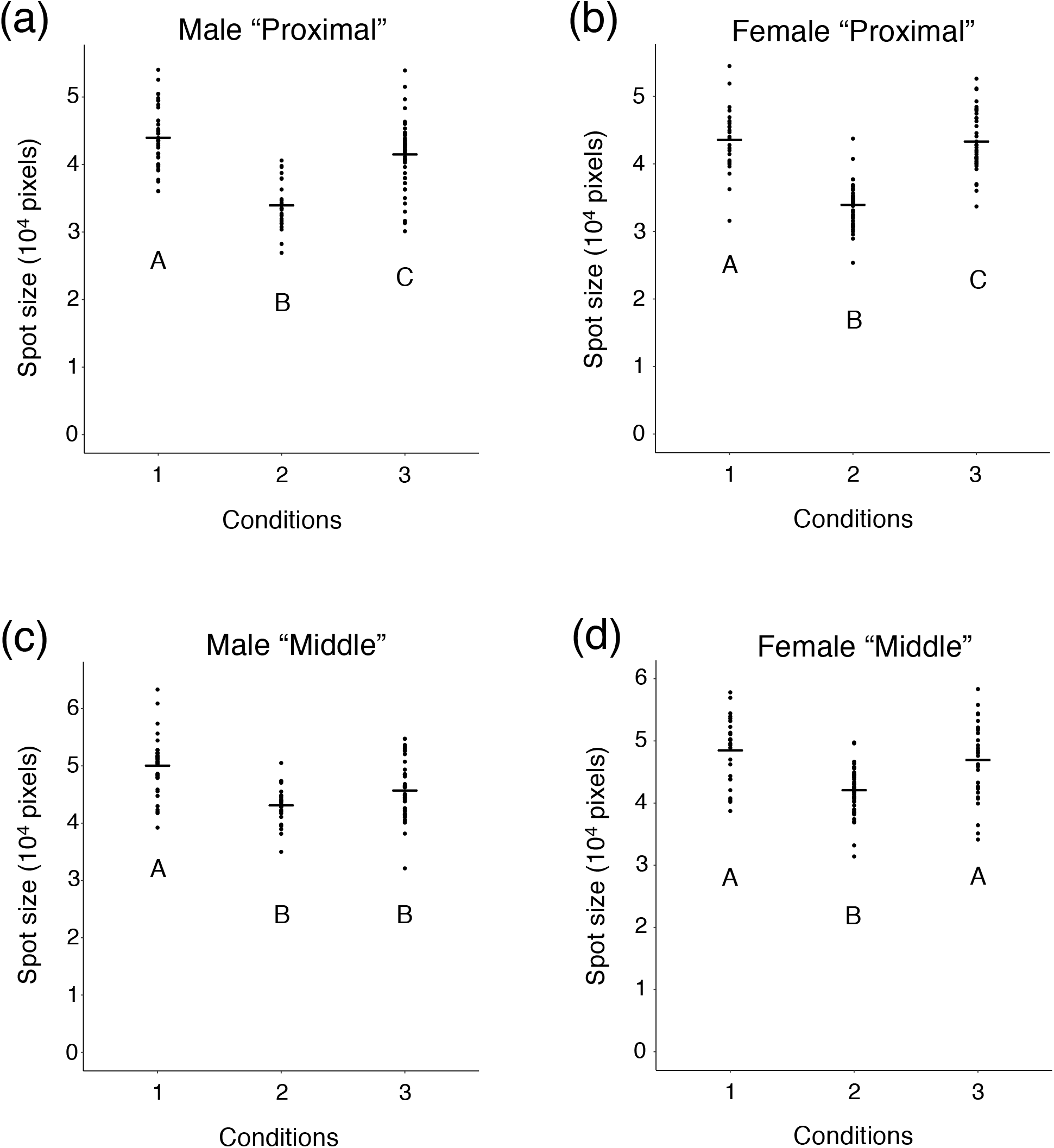
The spot size of wings from flies reared under “Condition 1”, “Condition 2”, and “Condition 3”. (a) Size of “Proximal” spots on wings from male flies. (b) Size of “Proximal” spots on wings from female flies. (c) Size of “Middle” spots on wings from male flies. (d) Size of “Middle” spots on wings from female flies. In all categories, there were significant differences between temperatures(*p* < 10^-6^, one-way ANOVA, degree of freedom = 2, *F* = 39.75 in (a), 78.34 in (b), 20.07 in (c), 17.77 in (d)). Different letters indicate significant differences (*p* < 0.05, Tukey’s HSD test). Black bars indicate mean values.

When the spot size is adjusted with wing size, the mean spot size of the wings from flies reared under “Condition 2” became the smallest in all categories (Figure 11). No significant difference between the spot size of flies reared under “Condition 1” and “Condition 3” was detected in any category (Figure 11).

**Figure 11.**
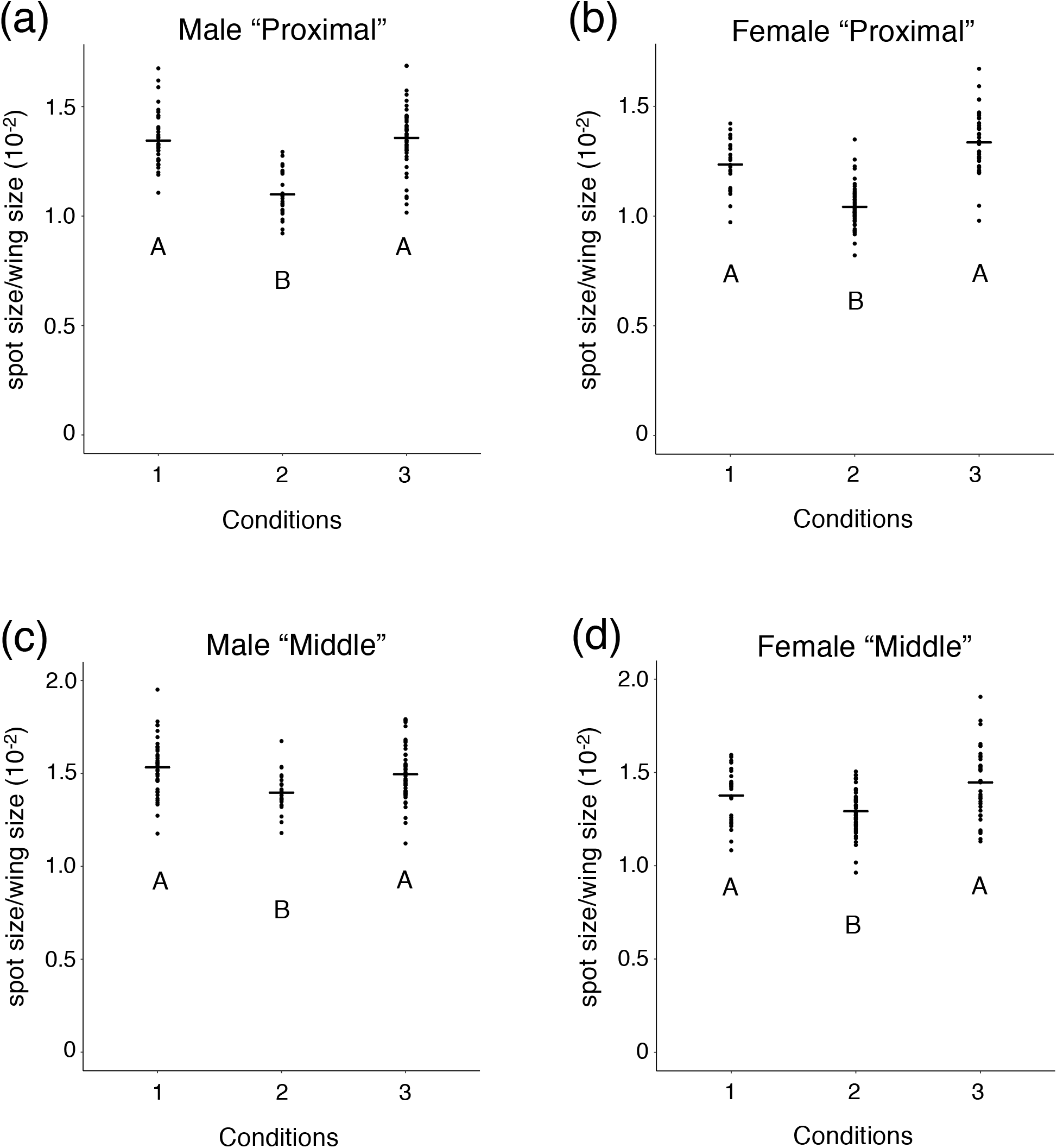
The spot size adjusted with wing size. Flies were reared under “Condition 1”, “Condition 2”, and “Condition 3”. (a) Size of “Proximal” spots adjusted with wing size from male flies. (b) Size of “Proximal” spots adjusted with wing size from female flies. (c) Size of “Middle” spots adjusted with wing size from male flies. (d) Size of “Middle” spots adjusted with wing size from female flies. In all categories, there were significant differences between temperatures (*p* < 10^-7^, Kruskal-Wallis rank sum test, degree of freedom = 2, *χ^2^* = 45.315 in (a), 62.827 in (b), 15.805 in (c), 16.804 in (d)). Different letters indicate significant differences (*p* < 0.05, Wilcoxon rank sum test with Bonferroni correction). Black bars indicate mean values.

Under “Condition 2”, “Proximal” spot was smaller than “Middle” spot (Figure 8b). Analyzing the ratio of “Proximal” size to “Middle” size, we found that the ratio becomes smaller under “Condition 2” both in males and females (Figure 12). A significant difference between the spot size of the flies reared under “Condition 1” and “Condition 3” was not observed (Figure 12).

**Figure 12.**
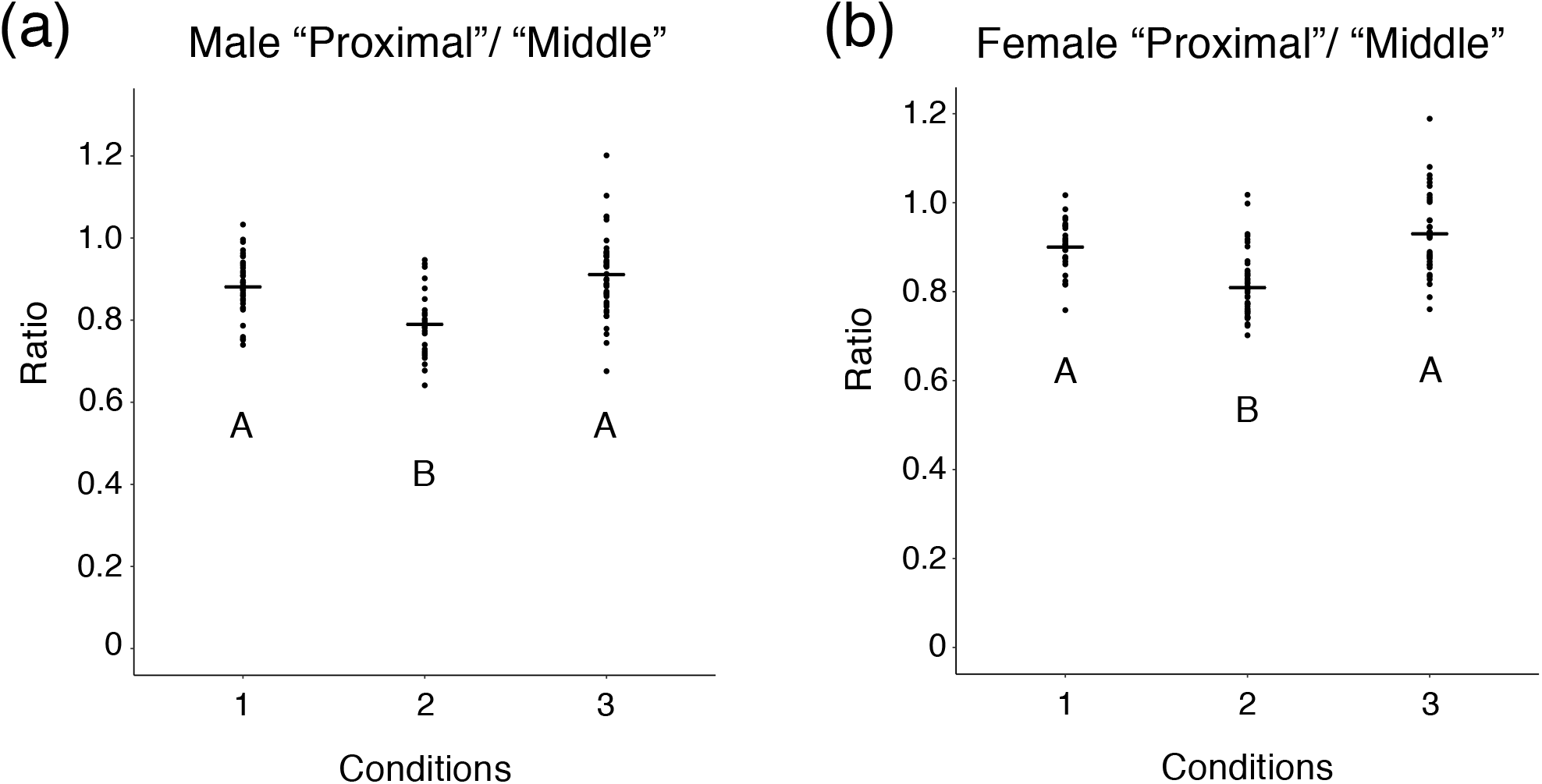
The ratio of “Proximal” spot size to “Middle” spot size. Flies were reared under “Condition 1”, “Condition 2”, and “Condition 3”. (a) The ratio in male flies. (b) The ratio in female flies. Both in males and females, there were significant differences between temperatures (*p* < 10^-5^, Kruskal-Wallis rank sum test, degree of freedom = 2, *χ^2^* = 26.414 in (a), 42.698 in (b)). Different letters indicate significant differences (*p* < 0.05, Wilcoxon rank sum test with Bonferroni correction). Black bars indicate mean values.

## 4 Discussion

In this study, we found that the wing size and the spot size of *D. guttifera* shows thermal plasticity. We also found that different spots have different reaction norms. The tendency of thermal plasticity we observed was almost identicle between males and females. Wing size becomes larger when flies are reared at lower temperatures (Figure 2, 3) as reported in other *Drosophila* species (Crill et al. 1996; Debat et al. 2003; Gilchrist and Huey 2004; Varón-González et al. 2020). As spot size itself changes between different temperatures (significant differences were observed by one-way ANOVA, Figure 4), the result suggests that spot size itself shows thermal plasticity. Differences in the ratio between the size of “Proximal” spots and the size of “Middle” spots depending on rearing temperature showed that “Proximal” spot and “Middle” spot have different reaction norms (Figure 7). When spot size is adjusted with wing size, the adjusted spot size became larger when the rearing temperature was higher (Figure 6). This might be because changes in wing size depending on temperature are much more drastic than changes in spot size itself (Figure 3, 4).

The results from experiments in which the rearing temperature is changed during the pupal period suggest that the thermal plasticity of wing size and that of spot size are independently regulated. For wing size, developmental stages before P4 (i) are the most sensitive period to the rearing temperature and there is no difference in sensitivity between pupal stages from P4 (i) to P14-15 and the stages after P14-15 (Figure 9). This result is similar to the tendency of wing size plasticity in *D. melanogaster* which shows that earlier developmental stages are more sensitive to the rearing temperature than later stages, such as pupal stages (French et al. 1998). Both the absolute spot size and the relative spot size adjusted with wing size becomes the smallest when flies are exposed to lower temperatures from P4 (i) to P14-15 (Figure 10, 11). Significant differences were detected between the spot size of the flies exposed to lower temperatures until P4 (i) and the spot size of flies exposed to lower temperature from P14-15 (Figure 10). However, there was no significant difference between them when spot size was adjusted with wing size (Figure 11). As a conspicuous phenotype, we observed that “Proximal” spot is smaller than “Middle” spot when flies were reared at lower temperatures (Figure 2a, b). This phenotype was observed only when flies were exposed to lower temperatures from P4 (i) to P14-15 (Figure 8, 12). The results for the spot size experiments showed that developmental stages sensitive to the rearing temperature are from P4 (i) to P14-15. The most sensitive stages for wing size and those for spot size are different. This suggests that the different developmental mechanisms produce thermal plasticity for wing size and spot size.

The relationship between the thermal plasticity of wing size and wing spot size is studied in another *Drosophila* species, *Drosophila suzukii*. In *D. suzukii*, male flies have a spot on a wing. Wing size and spot size change at different rearing temperatures, but spot size divided by wing size is almost robust (Varón-González et al. 2020). Although the genetic basis of spot formation in *D. suzukii* is unknown, the difference in tendencies of spot size changes in *D. guttifera* and *D. suzukii* might reflect differences in developmental mechanisms for spot formation.

Results obtained in this study suggest the possibility that the rearing temperature affects the mechanism of determining spot size by Wingless morphogen. Expression of *wingless* gene at campaniform sensilla starts at stage P6 (Werner et al. 2010) and the expression at presumptive spot regions can be detected at stage P12 (Fukutomi et al. 2021). After eclosion, epithelial cells at presumptive spot regions, which receive Wingless signaling, disappear (Fukutomi et al. 2017) and it can be considered that specification of pigmented regions by Wingless ends before eclosion. Therefore, stages at which Wingless specifies the spot regions are included in the period from P4 (i) to P14-15, which is sensitive to the rearing temperature in terms of determining spot size. If we assume that spot size is determined by diffusion of Wingless protein, spot size can change when the gradient of Wingless protein is altered. As our results show that exposure to 18 °C during the period including *wingless* expressing stages makes spot size smaller, the idea that the gradient of Wingless is altered at 18 °C is consistent with our results.

Factors other than the gradient of Wingless that can affect spot size should be also considered. One candidate phenomenon is the transportation of materials through wing veins. In *Drosophila* and *Ceratitis* species, the transportation of melanin precursors contributes to the proper formation of wing pigmentation after eclosion (True et al. 1999; Fukutomi et al. 2017; Pérez et al. 2018). However, our results suggest that the transportation of materials through veins after eclosion does not have a considerable effect on the thermal plasticity of wing spots. Exposure to 18 °C since pupal stage P14-15 did not produce a difference in spot size between “Proximal” spot and “Middle” spot, the conspicuous phenomenon that can be observed when flies were reared at lower temperatures (Figure 8, 12). As another factor, the possibility that a gene(s) other than *wingless* is (are) responsible for thermal plasticity can be considered. From the results that the adjusted size of “Proximal” spot and “Middle” spot showed different degrees of reductions in size at lower temperatures (Figure 6, 7, 11, 12), it is possible that a gene(s) which becomes differentially expressed between the two regions at lower temperatures contributes to the different degrees of size changes.

When our results are compared with the thermal plasticity of color patterns in butterflies, the comparison suggests that molecular mechanisms for the thermal plasticity of spots in *D. guttifera* might be different from that of color patterns in butterflies. *Bicyclus anynana*, a species of Nymphalid butterflies, has eyespots on the wing and the size of the “eyespot center”, the blue region located at the center of an eyespot, shows thermal plasticity (Prudic et al. 2011). The thermal plasticity of the “eyespot center” size is controlled by the change in titer of the molting hormone, 20-hydoxyecdysone (Riddiford 1993) and the most sensitive stage for thermal plasticity is the wandering stage before pupal formation (Monteiro et al. 2015; Bhardwaj et al. 2018). Although *wingless* gene is expressed at the presumptive “eyespot center” in the pupal period and is considered to have a role in pattern formation of eyespots (Özsu et al. 2017), the expression of *wingless* cannot be observed in wing discs of larvae and it can be observed on wings from pupal stages (Monteiro et al. 2006). The thermal plasticity of the “eyespot center” size might be controlled irrelevant of *wingless* gene.

Another Nymphalid butterfly, *Junonia coenia* shows thermal plasticity in wing color patterns by cold shock during the pupal period (Nijhout 1984). When heparin, a glycosaminoglycan that binds extracellular Wnt protein and affects the distribution of Wnt protein (Binari et al. 1997; Baeg et al. 2001), is injected into the pupae of *J. coenia*, the injected individuals will show a similar wing color pattern to the one induced by cold shock (Serfas and Carroll 2005). During the pupal period, *wingless* gene and *WntA* gene indeed are expressed in regions associated with some components of the wing color pattern (Martin and Reed 2010; Martin and Reed 2014). The necessity of *WntA* for color pattern formation is confirmed with CRISPR Cas9 system (Mazo-Vargas et al. 2017). Cold shock for *J. coenia* induces drastic changes that erase some components of the wing color pattern, and cold shock might affect the distribution of extracellular Wnt protein (Serfas and Carroll 2005). As loss of particular components of the wing color pattern is not observed when *D. guttifera* is exposed to a low temperature, molecular mechanisms for thermal plasticity of wing spots in *D. guttifera* might be different from that in *J. coenia*.

In conclusion, the size of wing spots around campaniform sensilla of *D. guttifera* shows thermal plasticity and reaction norms are different in different spots. The most sensitive period for the thermal plasticity of spot size includes the pupal stage at which *wingless* is expressed on wings. Our results suggest that the process for specifying the presumptive pigmented areas by Wingless is affected by temperature change, and they do not deny the possibility that genes other than *wingless* are responsible for the thermal plasticity. Our results also suggest that mechanisms for producing thermal plasticity in the wing spot size of *D. guttifera* might be different from mechanisms for thermal plasticity of color patterns in other insects. In the future, visualization of the distribution of Wingless protein (Panáková et al. 2005) in pupal wings of *D. guttifera* will help to understand the involvement of Wingless in the thermal plasticity. As there are other methods to transduce Wingless signaling than diffusion (Stapornwongkul and Vincent 2021), it will also help our understanding of how Wingless protein is allocated and specifies pigmented areas on wings in *D. guttifera*.

## Acknowledgements

We thank Sean B Carroll, and Thomas Werner for providing fly stocks; Tomohiro Yanone, Wataru Yamamoto, Namiho Saito, Hiroaki Osada, Machiko Teramoto and Tsuyoshi Katahata for fly stock maintenance; Koichiro Tamura, Masafumi Nozawa, Kentaro Tanaka, and Yige Luo for scientific advice; Takako Fijichika for creating scale bars in photo images. This work was supported by KAKENHI (21J00655) to Y.F.

## Author contributions

**Yuichi Fukutomi:** Conceptualization (lead); data Curation (lead); formal Analysis (lead); investigation (lead); methodology (lead); project administration (lead); resources (equal); validation (lead); writing – original draft preparation (lead); writing – review & editing (equal). **Aya Takahashi:** Conceptualization (supporting); resources (equal); supervision (equal); writing – review & editing (equal). **Shigeyuki Koshikawa:** Conceptualization (supporting); resources (equal); supervision (equal); writing – review & editing (equal).

## Data accessibility statement

Data obtained in this study are available from figshare at https://figshare.com/articles/dataset/Wing_size_and_wing_spot_size_of_Drosophila_guttifera/21760562

